# GRIMM-II: A Two-Stage Real-Time Algorithm for Nine-Locus HLA Imputation and Matching with Up to Three Mismatches

**DOI:** 10.64898/2026.03.28.715027

**Authors:** Ofek Kirshenboim, Amit Kabya, Regev Yehezkel-Imra, Yuli Tshuva, Martin Maiers, Loren Gragert, Pradeep Bashyal, Sapir Israeli, Yoram Louzoun

## Abstract

**Background:** The success of hematopoietic stem cell transplantation (HSCT) depends critically on human leukocyte antigen (HLA) matching between donor and recipient. While traditional matching focuses on five classical HLA loci (A, B, C, DRB1, DQB1), clinical practice increasingly considers extended typing at nine loci, including DPA1, DQA1, DPB1, and DRB3/4/5. Furthermore, emerging evidence supports transplantation with up to three HLA mismatches under post-transplant cyclophosphamide (PTCy) regimens. However, current donor search algorithms cannot efficiently identify donors with multiple mismatches across extended HLA loci in real-time.

**Methods:** We developed GRIMM-II (GRaph IMputation and Matching, version II), which comprises two novel algorithms: ML-GRIM (Multi-Locus GRIM) for HLA imputation across multiple loci, and ML-GRMA (Multi-Locus GRMA) for real-time donor-patient matching with up to three mismatches. Both algorithms employ a two-stage approach that combines efficient candidate reduction through graph-theoretic frameworks with detailed genotype comparison. ML-GRIM partitions genotypes into class I (HLA-A, B, C) and class II (remaining loci) components, enabling memory-efficient storage and rapid candidate identification. ML-GRMA searches a pre-imputed donor graph composed of donor genotypes and their sub-components, then computes asymmetric graft-versus-host (GvH) and host-versus-graft (HvG) mismatch probabilities to provide clinically relevant compatibility assessments. Both imputation and matching tools are available as a web application at https://grimmard.math.biu.ac.il/ and through GitHub repositories at https://github.com/nmdp-bioinformatics/py-graph-imputation (imputation) and https://github.com/nmdp-bioinformatics/py-graph-match (matching).

**Results:** We validated ML-GRMA and ML-GRIM using the WMDA3 (World Marrow Donor Association) validation dataset, successfully reproducing all previously reported matches while identifying numerous additional candidate donors not detected by previous algorithms. Further validation of ML-GRMA using 3,000 patients with artificially introduced mismatches (0-3 allele substitutions) demonstrated 100% sensitivity and specificity in identifying matching donors at expected mismatch levels. We validated ML-GRIM using simulated nine-locus typings derived from 8,078,224 US donors in the NMDP registry. The algorithm successfully imputed genotypes across variable numbers of typed loci while incorporating multiethnic haplotype frequencies. The algorithm achieved real-time performance with typical imputation times under one second and matching times of 1-13 seconds per patient for up to three mismatches, even when searching databases exceeding 8 million donors. Notably, ML-GRMA identified substantially more potentially suitable donors than traditional algorithms by accounting for the biological reality that GvH and HvG mismatches often differ, particularly for donors homozygous at specific loci. To evaluate ML-GRIM performance with low-resolution typing, we tested it on simulated 3-locus typings from the same population. The resulting imputation accuracy correlated with the mutual information between typed loci and complete genotypes.

**Conclusions:** GRIMM-II provides a scalable, memory-efficient solution for nine-locus HLA imputation and real-time identification of donors with up to three mismatches. The graph-based framework supports dynamic registry updates and can readily accommodate additional HLA loci and matching criteria as clinical knowledge evolves. By expanding the pool of acceptable donors while maintaining computational efficiency, GRIMM-II addresses a critical need in contemporary transplantation practice, particularly for patients from underrepresented ethnic minorities who face lower probabilities of finding perfectly matched donors.

## 1 Introduction

Hematopoietic stem cell transplantation (HSCT) remains a life-saving therapeutic intervention for patients with hematologic malignancies, immunodeficiencies, and various genetic disorders [13, 26]. The success of HSCT depends on the degree of human leukocyte antigen (HLA) compatibility between donor and recipient, as HLA mismatches can lead to graft-versus-host disease (GvHD), graft rejection, and increased transplant-related mortality [21, 11]. Traditionally, the gold standard for HSCT has been HLA-identical sibling donors or unrelated donors matched at high resolution for HLA-A, B, C, DRB1, and DQB1 loci (10/10 match) [20]. However, the probability of finding perfectly matched unrelated donors varies significantly across ethnic populations, ranging from approximately 75% for patients of European ancestry to less than 25% for those from underrepresented ethnic minorities [14, 8].

The landscape of HSCT has evolved considerably with advances in transplant conditioning regimens, GvHD prophylaxis, and supportive care [2, 29]. Contemporary clinical evidence demonstrates that carefully selected HLA-mismatched transplants can achieve outcomes comparable to fully matched transplants, particularly when advanced GvHD prevention strategies are employed [22, 27]. Studies have shown acceptable outcomes with single antigen mismatches [25, 28], and emerging data support the feasibility of transplants with up to three mismatches when appropriate patient and donor selection criteria are applied [6, 30]. Post-transplant cyclophosphamide (PTCy), T-cell replete haploidentical transplants, and other novel approaches have further expanded acceptable mismatch thresholds, making previously ineligible donors viable transplantation options [3, 12].

Although HLA matching in the PTCy context remains centered on four classical loci (HLA-A, -B, -C, and -DRB1), there is increasing interest in characterizing donor genotypes at DQA1, DQB1, DPA1, DPB1, and DRB3/4/5 (hereafter referred to as the nine loci) to avoid selecting donors who carry HLA antigens against which patients have pre-existing antibodies.

Among the additional HLA loci requiring consideration, DPB1 has emerged as particularly significant. Matching for both DPB1 alleles remains the optimal strategy to prevent acute GvHD, with studies demonstrating that DPB1 matching status significantly affects outcomes for recipients of unrelated donor stem cell transplants [30]. The complexity of DPB1 matching extends beyond simple allelic compatibility, involving sophisticated algorithms that account for T-cell epitope (TCE) groups and expression levels to predict clinical outcomes [7].

To support nine-locus HLA imputation and matching, we utilized population-specific nine-locus haplotype frequency estimates derived from large-scale registry data using a graph-based expectation-maximization (EM) framework. These frequencies extend classical five-locus HLA haplotypes (HLA-A, -B, -C, -DRB1, -DQB1) to include HLA-DPA1, -DPB1, -DQA1, and DRB3/4/5, enabling probabilistic modeling of extended class II diversity.

Current donor search algorithms have not adapted to this evolving landscape. Conventional matching algorithms employed by donor registries often implement rigid hierarchical matching strategies that may inadvertently exclude potentially suitable donors who fall outside traditional matching paradigms, as exemplified in our results. Moreover, most existing systems are optimized for identifying perfect or near-perfect matches. In contrast, we focus on identifying the large pool of mismatched donors with up to three mismatches. While solutions exist for extensive searches to find mismatched donors [9], or solutions to find the probability of multiple mismatches for donors that have up to 1 mismatch[4, 31, 10], there are no currently published identification capabilities for all donors with multiple mismatches. Finally, the dynamic nature of donor registry databases, with continuous additions and updates, necessitates algorithms capable of efficiently incorporating new data without requiring complete recomputation of matching results, in contrast to existing algorithms.

Extended HLA matching beyond the classical five loci presents unique immunogenetic and operational challenges. HLA-DQ and HLA-DP function as heterodimers formed from alpha and beta chains encoded by separate loci (DQA1/DQB1 and DPA1/DPB1), creating compatibility scenarios where gene-level matching may not reflect molecular compatibility [7]. DPB1 polymorphism particularly impacts acute GvHD risk through T-cell epitope (TCE) group classifications. Population diversity remains paramount, as haplotype frequencies vary dramatically—patients of European ancestry have 75% probability of finding 10/10 matched donors versus less than 25% for underrepresented minorities [14]. Clinical decision support must provide probabilistic compatibility assessments incorporating directional mismatch vectors (graft-versus-host and host-versus-graft) rather than symmetric scores [21]. DRB3/4/5 genes exhibit presence/absence variation across haplotypes, requiring algorithms to distinguish true null alleles from gene absence.

As such, nine-locus frequencies limit the applicability of existing imputation algorithms [9]. Classical imputation algorithms sequentially compare a list of possible haplotypes to identify all haplotype pairs consistent with a given HLA typing.

We previously developed a framework named GRIMM[18, 23, 16, 1] for graph-based imputation and matching. Imputation was performed by expanding all candidate phases and then checking all possible haplotypes in each phase using graph traversal. Matching was performed by comparing all imputed genotypes of the patient and donor using a graph traversal algorithm. The same framework was extended to multiethnic imputation and to an EM framework for producing haplotype frequencies [17].

However, neither nine-locus imputation nor matching can be applied with GRIMM at nine loci. Both imputation and matching require large imputation or matching graphs that cannot be maintained in memory using the current GRIMM formalism, as they require storing all sub-components of alleles or genotypes. To address the challenges of nine-locus imputation and matching, we propose adding a blocking step (rapid removal of candidate solutions) to GRIMM for both imputation and matching. This fundamentally reconceptualizes the donor search and imputation processes. In both cases, we propose highly memory- and CPU-efficient reduction of the candidate list (genotypes/donors) to a small subset using a graph-theoretic framework, followed by more standard genotype comparison. We denote this algorithm ML-GRIM (Multi-Locus GRaph IMputation) and ML-GRMA (Multi-Locus GRaph Matching).

The main novel aspects of the proposed algorithms are:

- A two-stage search with efficient candidate reduction followed by comparison of the reduced candidate list with the typing (in ML-GRIM) or the donors’ genotypes (in ML-GRMA).
- An efficient graph framework to maintain donors and haplotype graphs in memory.
- Asymmetric comparison of donor and patient genotypes, allowing for distinct graft-versus-host (GvH) and host-versus-graft (HvG) mismatch counts.
- Real-time search (typically 1 second per patient) with up to three mismatches.
- Real-time imputation (typically less than 1 second for nine loci per typing).

In the following Methods and Algorithm section, we present the details of ML-GRIM and ML-GRMA. In the Results section, we demonstrate the accuracy of ML-GRIM and ML-GRMA on simulated data. To clarify notation, we define GRIMM as the combination of GRIM and its matching algorithm, and GRIMM-II as the combination of ML-GRMA and ML-GRIM.

## 2 Methods and Algorithm

### **2.1** ML-GRIM Outline

ML-GRIM extends GRIM [16]. The input to ML-GRIM is an unphased and ambiguous genotype represented in genotype list string (GL-string) format [24], and the output is a list of the most probable genotypes in GL-string format along with their probabilities, based on the individual’s ethnicity information and population-specific HLA genotype frequency distributions. We considered the *k* most probable genotypes for each individual. In the WMDA3 analysis, we wanted to ensure all previously reported results are found and used *k* = 1, 000; in all other analyses, we used the more realistic *k* = 20.

As with ML-GRMA, ML-GRIM employs a two-stage approach: a blocking stage to limit candidates, followed by detailed comparison within a restricted candidate set. First, a subset of three loci is selected (for example, A, B, DRB1), and all other loci are temporarily ignored. Classical GRIM is then applied using only these three loci, and all genotypes consistent with the typing at these three loci are retained as the blocking stage output. For each of these genotypes (unphased and unambiguous), we verify consistency with the remaining typed loci, if any. If the initial typing contains fewer than three loci, only the available loci are selected, and classical GRIM is applied to those (Fig. 1).

**Figure 1:**
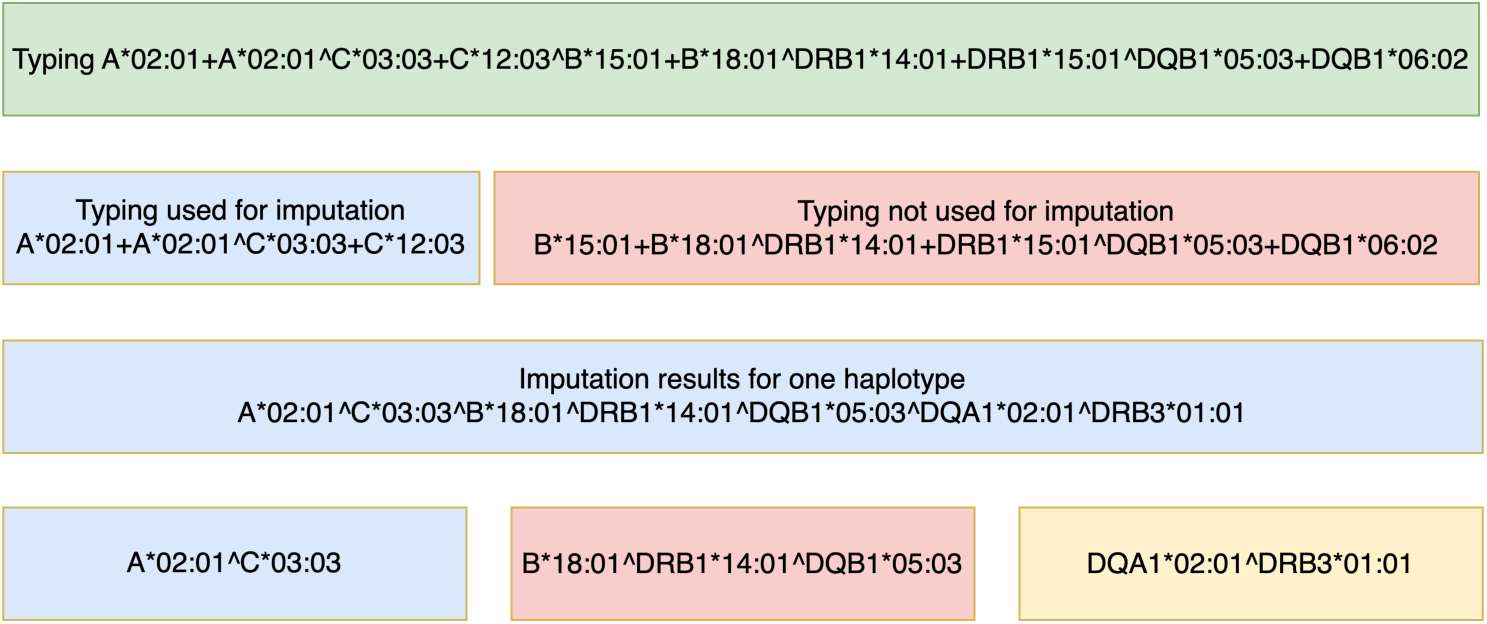
The input is an unphased and possibly ambiguous (some alleles are uncertain) or partial (not all alleles are typed) HLA genotype in GL-string format (top green box). In the blocking stage, a subset of loci (here: HLA-A and HLA-C) is selected for initial imputation (blue box), while other typed loci are temporarily ignored (red box). The classic GRIM algorithm is then applied to generate consistent haplotypes. The results (middle blue box) show a single imputed haplotype that combines the initial typing with additional inferred loci (e.g., DQA1, DRB3) based on population frequency priors. The bottom row demonstrates how the final imputed haplotype is constructed by integrating the blocking loci (blue), remaining typed loci (red), and fully imputed loci (yellow).

### **2.2** ML-GRMA Outline

We first discuss the general outline of the ML-GRMA algorithm, then detail each component (Fig. 2 upper plot). First, all donor HLA typings are imputed using ML-GRIM to produce a set of genotypes for each donor along with the probability of each genotype. The donors and their associated genotypes are represented as vertices in a graph, with edges connecting each donor ID to all high-resolution genotypes associated with it, weighted by genotype probability. ML-GRMA then constructs additional vertices for each genotype, representing the class I and class II genotype components (defined broadly; for example, a class I genotype comprises the pairs of A, B, and C alleles in a given genotype—six alleles total—while all other alleles constitute class II). Note that this is an arbitrary partitioning, and any grouping of loci into two approximately equal-sized groups can be used, since the blocking stage is only there to limit candidates and reduce computational cost. It has no effect on the final output. Each class I or class II genotype is connected to vertices representing all alleles it contains except one (e.g., two A alleles, two B alleles, and only one C allele). These are denoted subclass vertices. Subclass vertices are required to identify mismatched donors and patients in real time.

**Figure 2:**
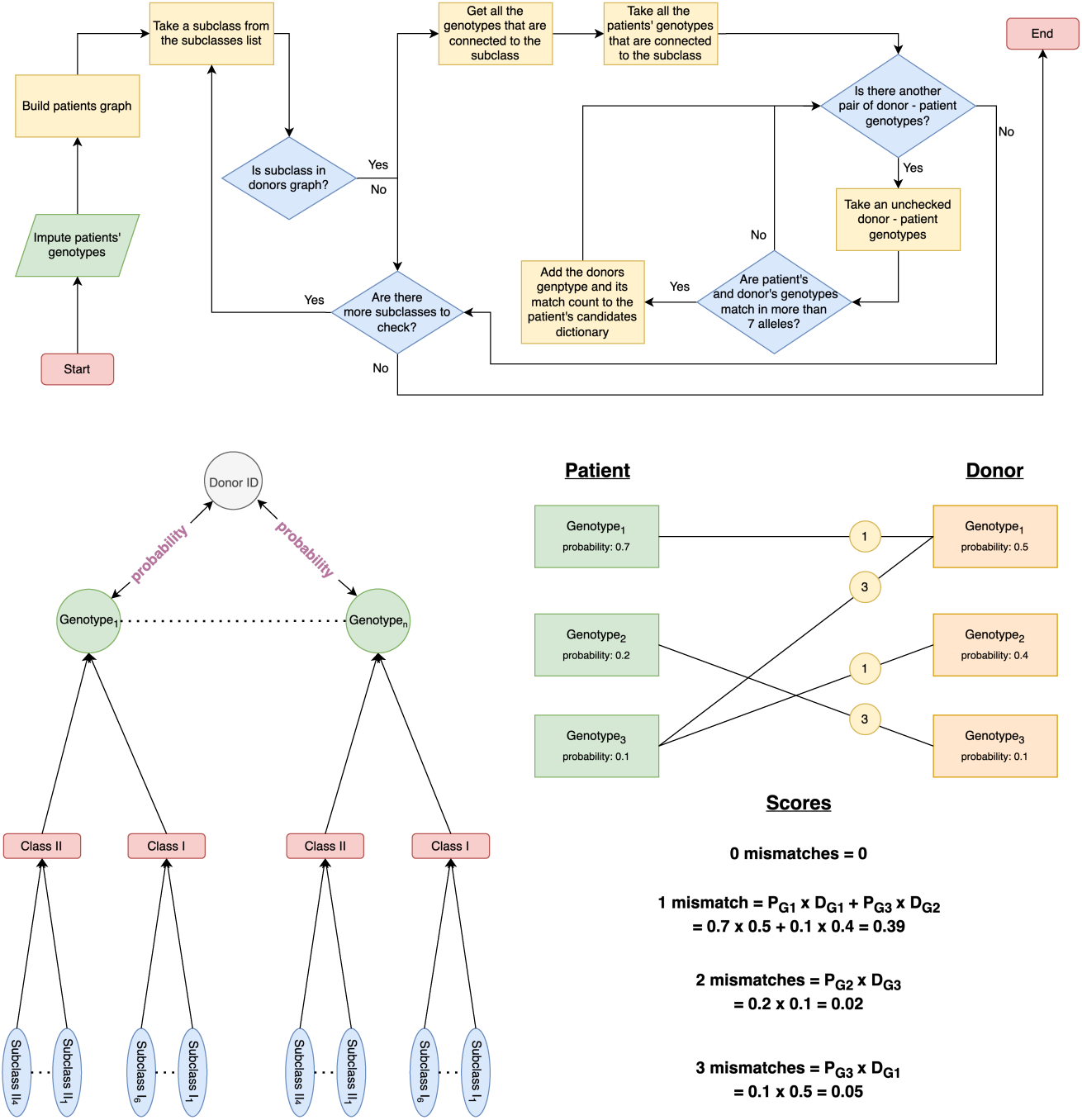
Schematic representation of the donor patient genotype matching workflow (ML-GRMA). The flowchart (top) describes the full algorithmic pipeline for processing patient genotypes, imputing missing data, constructing the possible high-resolution patient genotype graph, extracting subclasses, and systematically comparing each subclass to the donor genotype graph. For every donor patient genotype pair, the algorithm checks for no more than 3 mismatches (numbers in circles); valid pairs are recorded in the patient’s candidate dictionary together with their computed mismatch score. The bottom illustration depicts an example donor patient matching scenario, the structure of genotype connections, and the calculation of the number of mismatched alleles (0-3).

Given a new patient typing, nine-locus high-resolution unambiguous un-phased genotypes consistent with the typing and their probabilities are imputed using ML-GRIM. These genotypes are organized in a graph similar to the donor graph (Fig. 2 lower left). ML-GRMA then iterates over the patient’s subclass vertices (twice as many subclass vertices as the number of loci).

If a patient subclass vertex is found in the donor graph, all patient genotypes associated with this subclass are added to the genotype candidate list. The list of unique genotypes is then compared with patient genotypes using allele-by-allele comparison, retaining only results with at most three mismatches.

Finally, for each donor ID associated with genotypes in the genotype candidate list, we compute the match probability between the donor and patient (Fig. 2 lower right, and section Match probability). Only results exceeding a predefined probability cutoff are presented. In the WMDA3 analysis, a cutoff of 0 was used. In other comparisons, we used different cutoffs as detailed in Results.

### **2.3** Donor and Patient Graphs

The donor graph contains the list of possible unambiguous genotypes for all donors and their probabilities. All donor typings were imputed using ML-GRIM. The donor graph consists of four layers, each with different vertex types connected to the next/previous layer:

**IDs layer:** Donor IDs.
**Genotypes layer:** From each ID vertex, edges connect to appropriate genotype vertices with weights representing genotype probability for that donor typing (normalized to 1).
**Classes layer:** Each genotype is partitioned into two genotype classes (not haplotypes)—class I (HLA-A, -B, and -C; six alleles across two chromosomes) and class II (all remaining alleles; 12 alleles at nine loci or 4 alleles at five loci).
**Subclasses layer:** For each class, vertices are created by removing one allele, yielding, for example, five alleles for class I (Fig. 2). These subclasses enable identification of genotypes mismatched by up to three alleles.

The donor graph is maintained in a memory-efficient data structure. First, all alleles in donor genotypes are tallied, and each allele is represented by its serial number. In the matching context, null alleles and the absence of DRB3/4/5 are represented by the same value; the specific null allele is recovered during detailed comparison. The relationship between alleles and their serial numbers is maintained through a bidirectional dictionary. When comparing allele matches, serial numbers are actually compared. Other vertices are maintained through a hash function to a representative integer. The patient graph is created similarly to the donor graph, with a separate graph constructed for each patient.

### **2.4** Donor Candidates

If a genotype, class, or subclass vertex from the patient graph exists in the donor graph, ML-GRMA retrieves all genotypes with a path from this vertex as candidates. It then counts matches between these genotypes and the patient genotype. Only genotypes with at most three mismatches are retained. For each such genotype, ML-GRMA maintains the donors possessing that genotype and their match probabilities in a dictionary. The mismatch count is defined as the sum over loci of the maximum of GvH and HvG mismatches across the entire genotype. Note that this differs from most current algorithms that compute the maximum between GvH and HvG per locus before summing.

For GvH mismatches, we count the number of alleles present in the patient but absent in the donor. Similarly, HvG mismatches represent the number of alleles in the donor absent in the patient. In both cases, null alleles are not counted as mismatches. A mismatch is generally defined as the maximum between GvH and HvG mismatches. DRB3, 4, and 5 are considered a single locus, as no cases of more than one of these genes on the same chromosome have been reported. The absence of any allele at DRB3/4/5 is also counted as a null allele.

#### **2.4.1** Match Probability

For each matched donor-patient pair identified by ML-GRMA, the match probability between patient and donor is computed (Fig 2): **Definition:** Let *G* = {*g*_1_*, g*_2_*, . . ., g_k_*} and *P* = {*p*_1_*, p*_2_*, . . ., p_k_*} be the sets of patient genotypes and their probabilities, respectively. Let 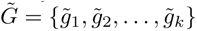 and 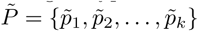 be the sets of donor genotypes and their probabilities, respectively. Let 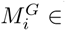 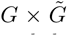 and 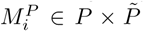 be the sets of matching genotype pairs for patient and donor with *i* matched alleles and their probabilities. The matching score between patient and donor is defined as:

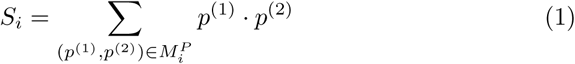

The analysis is performed separately for GvH, HvG, and the total number of mismatches.

#### **2.4.2** Donor Registry Data

We analyzed 8,078,224 US donors from the NMDP registry [15]. The dataset included donor CMV status, blood type, age, gender, race and ethnicity, registration year, and HLA typing.

#### **2.4.3** Simulated Typing

To evaluate imputation accuracy in a controlled setting, we generated simulated HLA typings from known nine-locus genotypes. First, phased nine-locus haplotype pairs were sampled according to the population-specific haplotype frequency distributions described above. These haplotype pairs were treated as ground-truth genotypes.

Simulated observed typings were then generated by masking loci to produce incomplete typings of varying resolution. Specifically, three-locus typings were created by retaining alleles at selected loci (e.g., HLA-A, -B, -DRB1) and removing all other loci. Typing ambiguity was introduced by collapsing alleles into multiple-allele codes or lower-resolution representations consistent with real-world registry typing practices. All simulated typings were encoded using GL-string format.

These simulated typings were then imputed using ML-GRIM, and imputation accuracy was assessed by determining whether the true underlying genotype was recovered among the top-k most probable imputed genotypes and by evaluating the posterior probability assigned to the true genotype.

#### **2.4.4** Simulated Mismatches

To validate mismatch detection, we randomly sampled 10,000 individuals from the cohort of 8,078,224. Each individual *j* had *r_j_* possible genotypes obtained through imputation. To ensure genotypes would not be missed due to low probability, we selected the highest posterior probability genotype for each individual as the reference sequence. We then generated artificial variants for each individual by performing *i*-substitutions, defined as random replacement of *i* alleles in the genotype sequence, where each substituted allele was replaced with a different possible allele at the same locus. We created 10 variants for each of *i* = 1, 2, 3 substitutions, resulting in 30 variants per individual. Sampled individuals were designated as patients, while their generated variants served as donors. For every patient-donor pair, we computed expected GvH, HvG, and mismatch scores to serve as ground truth labels. Since substitutions were performed randomly, in some cases a replaced allele could coincidentally match the other allele at the same locus, leading to no effective change in corresponding compatibility metrics.

For validation, we constructed input files where original sampled individuals were listed as patients, and the donor pool consisted of both their artificially generated variants and the remaining cohort (i.e., all individuals except those designated as patients). ML-GRMA was then executed on these inputs to compute compatibility scores for all patient-donor pairs. Algorithm output was compared against precomputed ground truth labels, confirming that ML-GRMA correctly identified all donor variants and reproduced expected GvH, HvG, and mismatch values.

The version of the code used for all the results here is the version available at the githubs of ML-GRMA and ML-GRIM at https://github.com/nmdp-bioinformatics/py-graph-match and https://github.com/nmdp-bioinformatics/py-graph-imputation

#### **2.4.5** WMDA Matching Validation Dataset 3

To validate ML-GRIM and ML-GRMA against an established international benchmark, we used the WMDA Matching Validation Task 3 (WMDA3) dataset, as defined by Bochtler et al. [5]. This dataset was developed by the World Marrow Donor Association (WMDA) to provide a standardized reference for evaluating probabilistic HLA matching algorithms.

The WMDA3 dataset consists of 1,000 simulated patients and 10,000 simulated donors, yielding 10 million patient-donor pairs. Genotypes were generated by independently sampling two-phased haplotypes from high-resolution U.S. White five-locus(HLA-ÃC~B~DRB1~DQB1) haplotype frequencies. Observed typings were then derived by introducing realistic patterns of typing ambiguity and missing data to mimic registry conditions. All typings are represented using GL-string syntax and include multiple-allele codes, ARD-grouped alleles, and untyped loci.

The WMDA3 task requires matching algorithms to compute allele-level match probabilities through imputation using high-resolution haplotype frequency data and without trimming the space of possible diplotypes. Importantly, this dataset explicitly penalizes algorithms that discard low-probability haplotype pairs, making it particularly suitable for validating the correctness of probabilistic matching frameworks.

For validation, both patients and donors were imputed using ML-GRIM with the provided five-locus haplotype frequencies. ML-GRMA was then applied to compute match probabilities and mismatch classifications. Results were compared against the WMDA3 reference outputs, confirming that GRIMM-II reproduces all previously reported matches and probabilities to within rounding error, while additionally identifying valid higher-mismatch matches not reported by traditional algorithms due to trimming or restrictive mismatch definitions.

## **3** Results

### **3.1** ML-GRIM

We previously developed GRIMM, a framework for HLA imputation and donor-patient matching. The imputation is based on expanding all candidate phases of the typing, which requires 2^Number^ ^of^ ^phases−1^ operations, and maintaining in memory all candidate haplotype sub-components. This is feasible with up to six loci but exceeds current machine memory capacity for nine loci. Beyond memory cost, the computational cost of expanding all possible phases with nine-locus typing is prohibitive. Similarly, matching requires maintaining all donors’ possible genotype sub-components (e.g., for a 10-allele (5-locus) genotype, vertices for all 9-, 8-, 7-allele sub-components), which is infeasible for an 18-allele (9-locus) genotype.

To address this, we propose two-stage approaches for both imputation and matching. A first blocking stage identifies all candidates (with no false negatives—not missing any genotype—but many false positives—candidate genotypes inconsistent with the typing). The second stage is less efficient but applies to a small number of candidates and removes all false positives, yielding a fully accurate solution for both imputation and matching.

For imputation, given typing composed of any number of input loci: if the typing has more than three loci, only the three most polymorphic loci (as defined by number of alleles in the frequencies) in the typing are selected (e.g., A, B, DRB1). We denote the full typing of patient *i* as 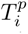 and the short typing with only three loci or fewer as 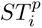 Typed loci not in 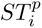 are denoted as the remainder, *R_i_*. We then perform classical GRIM imputation using only 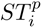 as input. Note that this requires maintaining only three-locus combinations in memory and expanding only four phases (2^3−1^). The resulting list of genotypes and haplotype pairs consistent with the three-locus typing are denoted *SG_i_*(*j*) and *SH_i_*(*j*), respectively. These may not be consistent with *R_i_*. The difference between *SG_i_*(*j*) and *SH_i_*(*j*) is that *SG_i_*(*j*) ignores haplotype phasing.

We then compare each genotype/haplotype pair *j* in *SG_i_*(*j*)*/SH_i_*(*j*) with the remaining typing *R_i_*. We compare only loci in *SG_i_*(*j*)*/SH_i_*(*j*) that are included in *R_i_*. Comparison is performed by checking whether *SG_i_*(*j*)*/SH_i_*(*j*) is included in *R_i_* and consistent with the chromosomal division of *R_i_* using a dictionary (hash table). For example, assume typing with A, B, C, DRB1, and DQB1. We partition the typing into 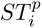 composed of A, B, and C, and *R_i_*. We then use 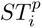 (i.e., the A, B, and C loci) as input to classical GRIM, obtaining resulting genotypes with nine loci. However, since only DRB1 and DQB1 are included in *R_i_*, ML-GRMA uses only these loci in *SG_i_*(*j*) and *SH_i_*(*j*). Assume, for example, *R_i_*= DRB1^∗^15 : 03*/*DRB1^∗^15 : 01^DRB1^∗^07 : 01 + DRB1^∗^01 : 02^DQB1^∗^05 : 01, and we have two matching results: *SG_i_*(1) = DRB1^∗^15 : 03^DRB1^∗^15 : 01 + DRB1^∗^01 : 02^DQB1^∗^05 : 01 and *SG_i_*(2) = DRB1^∗^15 : 03^DRB1^∗^07 : 01 + DRB1^∗^01 : 02^DQB1^∗^05 : 01. While both are included in the typed alleles, only the second is consistent with the phasing of the typing and is therefore used. We then compute the probability of each genotype or haplotype pair as performed in GRIM (Fig. 1).

### **3.2** ML-GRMA

A similar approach is used for matching (Fig. 2). Directly comparing all donor genotypes consistent with any patient genotype is computationally prohibitive. Similarly, maintaining in memory a direct list of all donor genotypes and their sub-components (e.g., for a nine-locus genotype, all sub-components with fewer than nine loci) is also prohibitively expensive. Instead of these extreme options—serial comparison (high CPU) or graph traversal (high memory)—limited information on each donor genotype is used to drastically reduce the number of candidate donor genotypes through graph traversal. This limited set of donors is then compared with the patient serially.

To find all candidate genotypes with up to three mismatches between donor and patient, we first pre-impute all donors using ML-GRIM. We partition each genotype into two groups of alleles (denoted here for simplicity as class I and class II, though any partition can be used). For example, assume a 5-locus genotype with 10 alleles: *A*_1_*, A*_2_*, B*_1_*, B*_2_*, C*_1_*, C*_2_, DRB_1_, DRB_2_, DQB_1_, and DQB_2_ (these are allele positions in the genotype, not allele names). ML-GRMA partitions these into class I (*A*_1_*, A*_2_*, B*_1_*, B*_2_*, C*_1_*, C*_2_) and class II (DRB_1_, DRB_2_, DQB_1_, DQB_2_).

If a patient and donor genotype have at most three mismatches, then either their class I or class II has at most one mismatch. Suppose, without loss of generality, that the donor and patient class II have at most one mismatch; then either “DRB_1_, DRB_2_, DQB_1_”, “DRB_1_, DRB_2_, DQB_2_”, “DRB_2_, DQB_1_, DQB_2_”, or “DRB_1_, DQB_1_, DQB_2_” are fully equivalent in the patient and donor genotype. These four combinations are denoted CIIM_1_ (class II minus 1 allele). For a *K*-locus genotype, there are *K* CIM_1_ and CIIM_1_ combinations. ML-GRMA produces all these combinations for each donor genotype. G iven a patient, ML-GRMA imputes candidate genotypes using ML-GRIM, then expands all class I and class II genotypes and all CIM_1_ and CIIM_1_. If the donor and patient share at least one CIM_1_ or CIIM_1_ vertex, the donor is a candidate. ML-GRMA then iterates over candidates and directly compares their full imputed genotypes to verify at most three mismatches.

In contrast to previous methods, ML-GRMA computes GvH and HvG mismatches separately. GvH mismatches represent the number of alleles present in the patient but absent in the donor, and vice versa for HvG. In both cases, null alleles are not counted as mismatches. The number of mismatches is the maximum of GvH and HvG across the entire genotype.

This definition of total mismatches as “max(total GvH, total HvG)” differs from accepted registry and clinical conventions. We define this as a new compatibility metric that is less conservative than current definitions. However, it accounts for the biological reality that a mismatch is either GvH or HvG, and that counting each locus separately is not biologically realistic.

### **3.3** ML-GRIM and ML-GRMA Validation

To test algorithm accuracy, we developed a set of simulations and validation measures. All ML-GRIM validations were performed against simulated typings where the ground truth was known. ML-GRMA validation was performed against simulated mismatches based on real typings. The following validations were performed (see Appendix, Table 1 for summary):

**ML-GRIM and ML-GRMA validation on WMDA3:** We compared the performance of combined ML-GRIM and ML-GRMA on a set of 5-locus simulated patients used for previous algorithms [5]. First, ML-GRIM was applied to patient and donor typings using simulated single-population haplotype frequencies. We then verified that the numbers and probabilities of full matches (10/10) or single mismatches (9/10) were consistent with published results.
**ML-GRIM validation using 9-locus simulated typing:** We simulated 3-locus typings from generated 9-locus genotypes and compared the imputation accuracy of ML-GRIM with the previously validated version of GRIM. For comparison, we excluded typings or frequencies with DPA1, DRB3/4/5, or DQA1 to ensure compatibility with previous algorithms. We then repeated the analysis using all loci combinations for 3-locus typing and 9-locus frequencies, comparing results to theoretical computations.
**ML-GRMA validation:** For GRMA validation, we used 8 million donor typings from the US NMDP registry and imputed their genotypes. We then selected 3,000 donors and designated them as patients. For each patient, we added 10 copies with either 0, 1, 2, or 3 alleles randomly replaced by another allele in their most likely genotype to the donor list. We added all these “mutated donors” to the donor list and searched the 3,000 patients in the enlarged donor list using ML-GRMA.

### **3.4** WMDA3 Validation of ML-GRIM and ML-GRMA

The WMDA3 dataset contains 10,000 simulated donors and 1,000 patients with appropriate single-population frequencies. We imputed each patient using ML-GRIM and performed matching between donors and patients using ML-GRMA. All reported matches were detected using the combination of ML-GRMA and ML-GRIM. The 9- and 10-match rate probabilities assigned by ML-GRIM and the original direct comparison are identical. Differences were at most 1%, resulting from rounding errors reported in the original results (GRMA9 vs. WMDA9 and GRMA10 vs. WMDA10 in Fig. 3, upper row). Typical imputation time is much less than one second. Typical matching time is 1-2 seconds (Fig. 3, second row). However, a large number of matches not previously reported were identified using ML-GRIM and ML-GRMA, mainly because no trimming of low-frequency results was performed in ML-GRIM and ML-GRMA. Note that ML-GRMA also reports 2- and 3-mismatch results, in contrast to existing algorithms (Fig. 3, bottom row).

**Figure 3:**
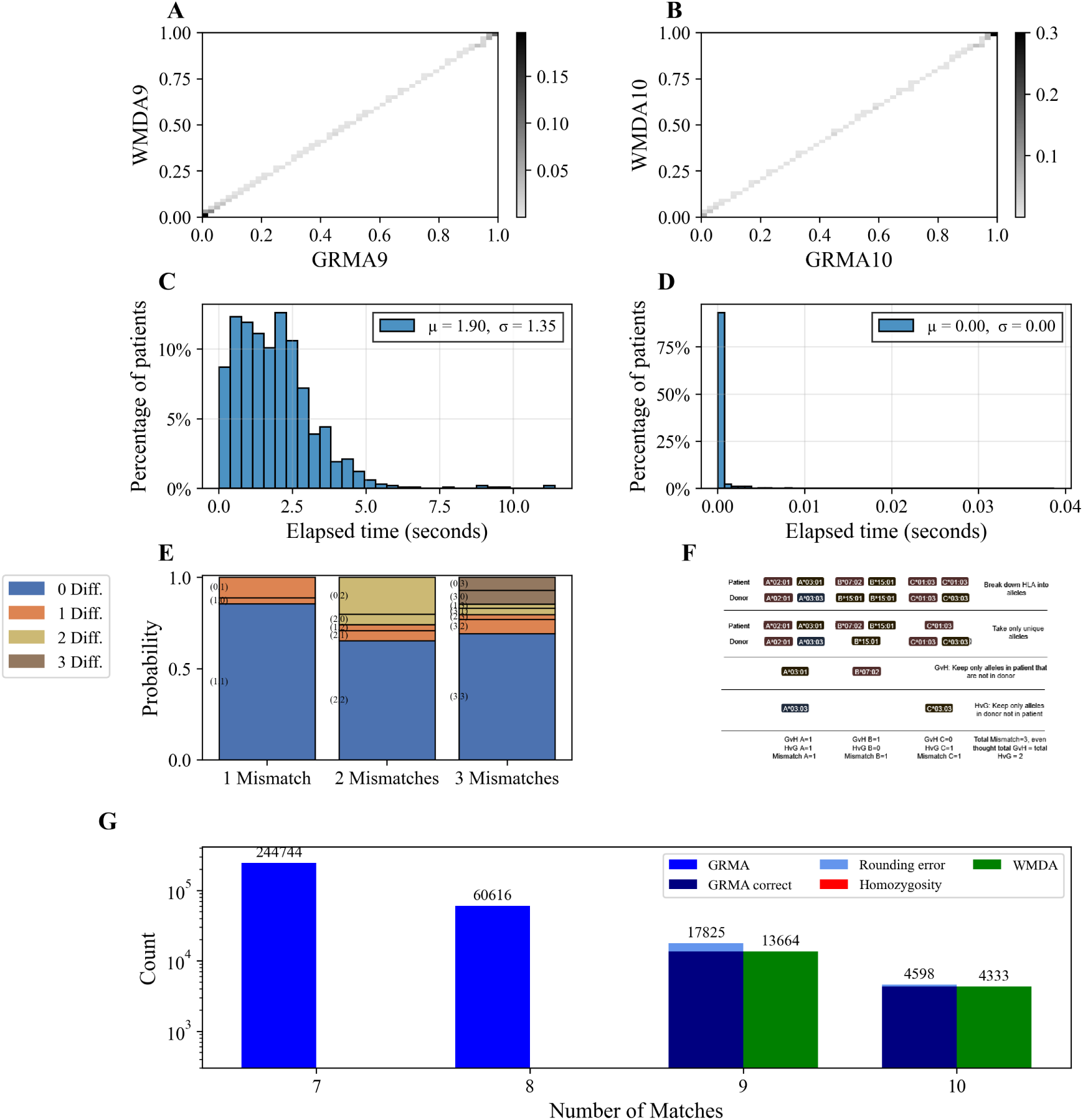
WMDA3 Validation. All the results below relate to the WMDA3 analysis. Subplots A and B. comparison of probability assigned to matches in ML-GRMA and in the direct enumeration (WMDA). The probabilities are the same (up to a rounding error of less than 1%). WMDA9 and 10 represent the 9 and 10 alleles matches from WMDA3, while GRMA9 and GRMA10 relate to ML-GRMA. Subplots C and D - runtimes in seconds of (C) matching and (D) imputation. The typical matching time is 1 second, while the imputation time is approximately 0.01 seconds. E - fraction of cases where the directional (GvH and HvG) mismatches are different from the total mismatches; blue represents equal directional and bidirectional mismatches, orange/beige and brown represent differences of 1,2, and 3 mismatches between the directional and undirectional mismatches, respectively. F. Schematic description of a case where the current definition would declare 2 mismatches, while the max of GvH and HvG would produce a single mismatch. G. Comparison of matched donor number between the reported WMDA3 and ML-GRMA. ML-GRMA finds more 0 and 1 mismatched donors.

An important difference between ML-GRMA’s match definition and existing algorithms is the treatment of homozygotes. We define the number of mismatches as the maximum between the number of mismatches from donor to patient (defined as GvH) and from patient to donor (defined as HvG) (Fig. 3, third row). Classical algorithms measure this maximum per locus and then sum across loci. As such, if there is a single GvH mismatch at one locus and a single HvG mismatch at another locus, existing algorithms would define that as two mismatches. We favor this new definition since in practice, both GvH and HvG across the entire genotype are 1; thus, there is never more than 1 mismatch in these cases. Note that differences between HvG and GvH are very frequent and typically represent 30% of cases with 2 and 3 mismatches (Fig. 3, bottom row). Since the algorithms are deterministic, there are obviously no false positives.

As mentioned, this definition differs from current clinical practice, and as such, the effect of using either the here proposed mismatch definition or the classical mismatch definition on the outcome of solid organ transplants and HSCT remains to be tested. The ML-GRMA code proposes reports both classical and current definitions of mismatches to allow for such comparisons.

### **3.5** Simulated 3-Locus Typings

Given that ML-GRIM can be applied to 9-locus typing but classical GRIM cannot, we first tested all combinations of A, B, C, DRB1, and DQB1 and assessed their imputation accuracy in the old (GRIM) and new (ML-GRIM) versions on 5- and 6-locus frequencies. The 5- and 6-locus frequencies were obtained from 9-locus frequencies by summing all haplotype frequencies sharing the same 5 or 6 loci, respectively.

We reported the first 20 most frequent imputed genotypes and tested the fraction of cases where real genotypes were found within these 20 imputed genotypes. We compared these fractions for GRIM and ML-GRIM for 5 and 6 loci, and for 9 loci for ML-GRIM, testing whether the real genotype (i.e., the genotype used for simulation) was recovered. The fraction of cases where the real genotype was found within the first 20 results for 5 and 6 loci is identical for GRIM and ML-GRIM. However, when comparing this number between 5, 6, and 9 loci, the fraction of cases where the real genotype is found is more than twice as high for 5 loci as for 9 loci (Fig. 4A). We then computed, for cases where real haplotypes are included in the imputed genotype, the total probability assigned to the real haplotype pair. Results are again similar for GRIM and ML-GRIM, and the average probability assigned to the real solution is twice as high in the 5-locus case as in the 9-locus case, as expected from the much larger number of options for 9 loci (data not shown).

**Figure 4:**
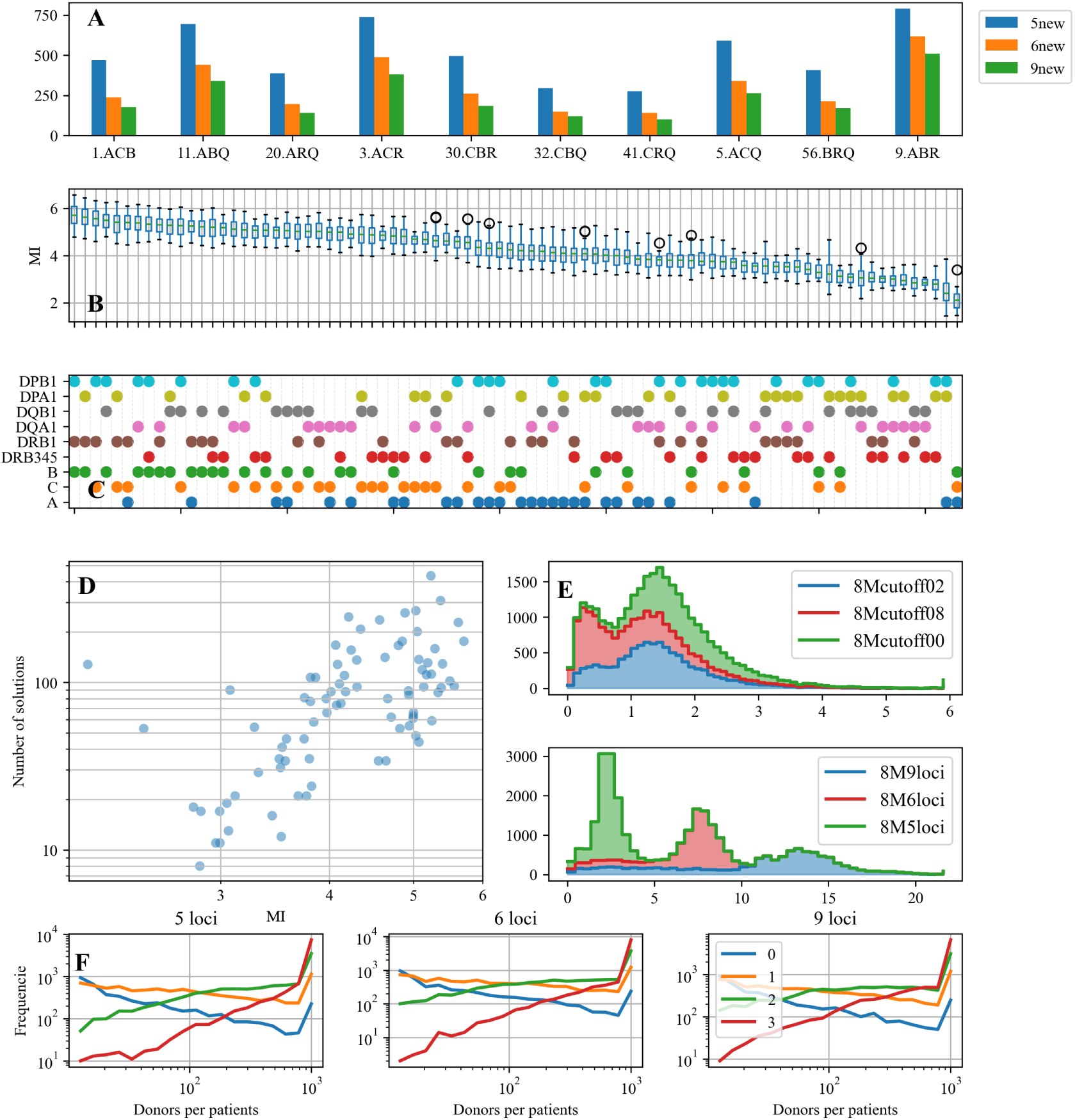
**Simulated typing validation**. A - number of cases where the simulated genotype was found in the imputed results of simulated typing of 3 loci, as a function of the loci used (out of 3,000 simulated typing), and the number of loci in the frequencies (different colors of bars). B - boxplots of mutual information (MI) between each pair of 3 loci and the other 6 loci. We computed the MI for each of the 21 detailed populations and 5 broad populations. The width of the distribution (boxes represent 25th-75th percentile) is narrow compared to the difference between combinations of 3-loci. C - The locus included in each combination in B. D - relation between MI and number of solutions found. As expected, when the MI is high, the number of solutions found is high. E - run time as a function of the cutoff used (using 5 loci) and of the number of loci used (using a 0 cutoff). F - histogram of the number of matching donors found with 0-3 mismatches, for 5,6 and 9 loci).

To understand this distribution, we computed the mutual information between frequencies in a set of loci and frequencies of all remaining loci for all populations studied. Mutual information is consistent among populations (Fig. 4B, C) and indeed explains most variability in the fraction of cases where the real genotype was found (Fig. 4D)

### **3.6** GRMA Validation

For ML-GRMA validation, we selected 3,000 donors and designated them as patients (while also retaining the original donor in the donor pool). We then introduced 1, 2, and 3 allele replacements by randomly swapping an allele with another. We added these “mutated” genotypes to a set of 8 million donors imputed to either 5, 6, or 9 loci. We computed whether these mutated donors were identified at 0, 1, 2, or 3 mismatches. All donors, without exception, were found with their appropriate expected number of mismatches. The time required to find a typical donor was 2.7, 9 and 13 seconds for 5, 6, and 9 loci, respectively, on a standard Intel i7 CPU.

All donor candidates are first tallied, and only then is a cutoff on frequencies applied. As such, while the cutoff can affect the number of candidate donors, it does not affect search time. To demonstrate this, we used three cutoffs on match probability between donor and patient (0%, 20%, and 80%) and computed run time for 5-locus matching (Fig. 4E), with no difference between cutoffs.

We limit the search to three mismatches. To verify that three mismatches are sufficient, we computed for the same simulated patients the number of donors with 0, 1, 2, and 3 mismatches at a probability greater than 0 (Fig. 4F). Interestingly, the number of 0- and 1-mismatch donors is low for most patients at either 5, 6, or 9 loci. However, the number of 3-mismatch donors is large, and most patients reached the threshold of 1,000 3-mismatch donors. Only 10 out of 10,000 simulated patients lacked a 3-mismatch donor.

## 4 Visualization and Web Server Interface

To enable practical use of ML-GRIM, we developed a web server application that combines a tool translating different typing input formats to standard GL-strings, denoted pyard (available at https://github.com/nmdp-bioinformatics/py-ard), with ML-GRIM. The combined tool is named GRIMMARD and is available as a GitHub repository (https://github.com/nmdp-bioinformatics/grimmard/) or as a web server with US and Israeli frequencies (https://grimmard.math.biu.ac.il/).

GRIMMARD receives as input: (A) the populations of interest, (B) the loci of interest, and (C) typing introduced as a GL-string or as allele numbers/codes at different loci. GRIMMARD allows mixed input combining serology and genetic typing at different resolutions (Fig. 5A). The outputs of GRIMMARD are:

1. unphased genotypes and their probabilities, (2) phased pairs of haplotypes and their probabilities, and (3) the probability of each candidate haplotype with its probability in each population of interest (Fig. 5B-D). Typical GRIMMARD run time is less than one second.

**Figure 5:**
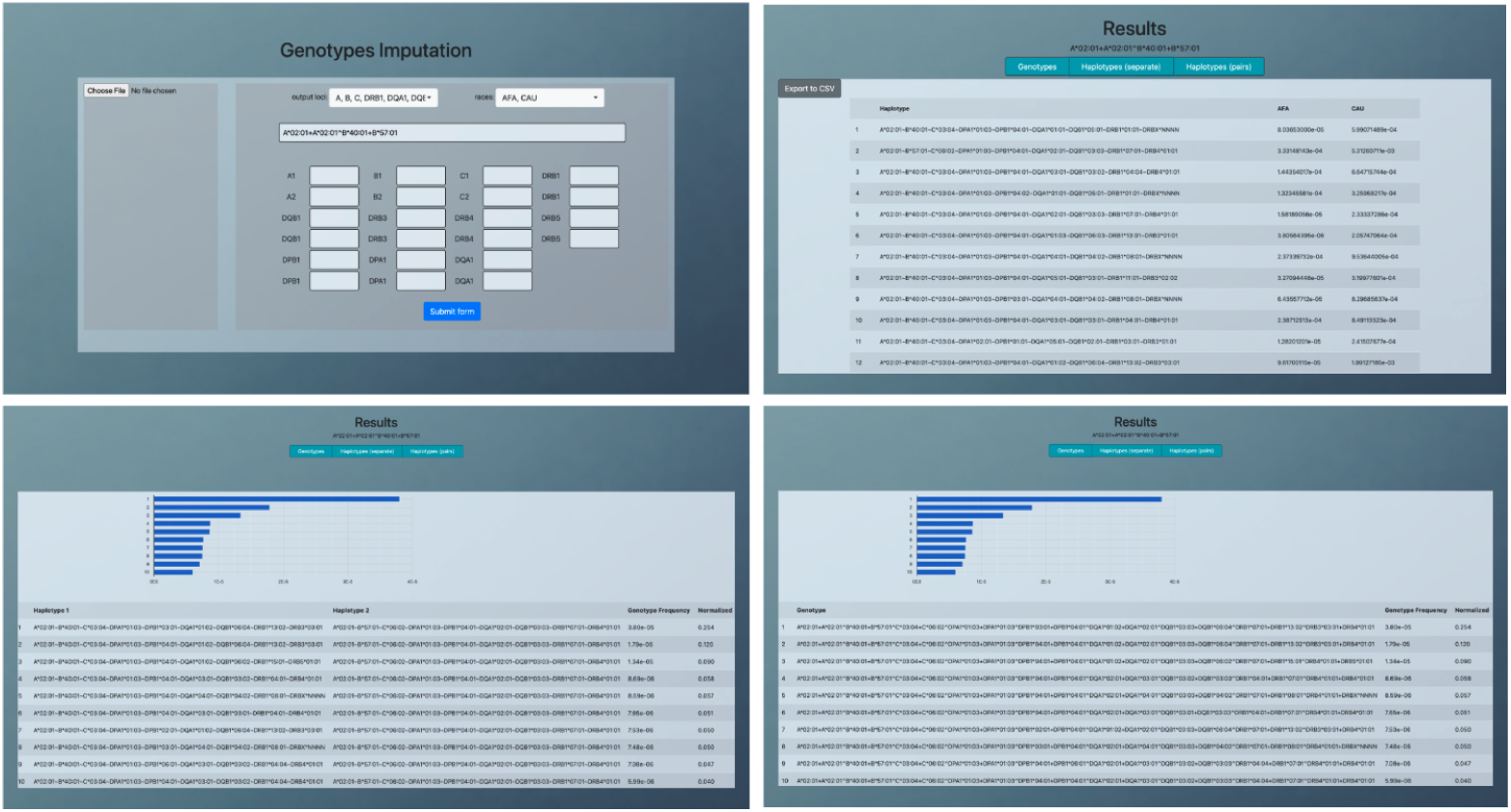
GRIMMARD web-server interface. Upper left: Input form accepting mixed-resolution HLA typing data in various formats, including GL-strings, allele codes, or serology. Upper right: Ranked unphased genotypes with their probabilities and frequencies. Lower left: Phased haplotype pairs with associated probabilities. Lower right: Individual haplotype probabilities across selected populations.

## 5 Discussion

The landscape of hematopoietic stem cell transplantation has evolved dramatically over the past two decades. Advances in conditioning regimens, GvHD prophylaxis strategies including post-transplant cyclophosphamide [22, 3], and improved supportive care [2, 13] have fundamentally altered the acceptable boundaries of HLA matching. Contemporary clinical evidence demonstrates that carefully selected HLA-mismatched transplants can achieve outcomes comparable to fully matched transplants, particularly when modern GvHD prevention strategies are employed [27]. However, donor search algorithms have not kept pace with these clinical advances, often employing rigid hierarchical matching strategies optimized for identifying perfect matches rather than systematically exploring the expanded space of clinically acceptable mismatched donors.

GRIMM-II addresses this gap by providing the first real-time algorithm capable of efficiently identifying donors with up to three mismatches across nine HLA loci. The twostage approach, combining graphtheoretic candidate reduction with detailed genotype comparison, overcomes the computational and memory constraints that have previously limited multilocus matching algorithms [23, 17]. Our validation demonstrates that the algorithm maintains perfect accuracy while achieving computational performance suitable for clinical implementation, with typical search times of 1-2 seconds per patient for 5 loci and up to 15 seconds for 9 loci, even when querying databases exceeding 8 million donors.

The extension from five-locus to nine-locus matching reflects evolving clinical understanding of factors influencing transplant outcomes. While HLA-A, B, C, DRB1, and DQB1 have traditionally formed the core of donor selection [21, 20], accumulating evidence supports the clinical importance of additional loci. DPB1 matching, in particular, has emerged as significantly impacting acute GvHD risk, with studies demonstrating that matching for both DPB1 alleles provides optimal protection [30, 28]. The complexity extends beyond simple allelic compatibility to encompass T-cell epitope groups and expression level matching algorithms that predict immunological consequences [7].

Similarly, HLA-DQA1 matching has clinical significance related to the unique biology of HLA-DQ molecules. These heterodimers, formed from *α* and *β* chains encoded by separate loci, create matching scenarios where a single DQB1 mismatch can result in one, two, or three mismatched DQ molecules, depending on DQA1 compatibility [19]. The inclusion of DRB3/4/5 loci, while technically complex given their variable presence across haplotypes, provides additional immunogenetic information relevant to transplant outcomes. GRIMM-II’s framework accommodates this complexity by treating the absence of these loci equivalently to null alleles, enabling consistent matching logic across diverse genetic backgrounds.

A distinguishing feature of ML-GRMA is its separate computation of GvH and HvG mismatches, reflecting the biological reality that these represent distinct immunological challenges. GvH mismatches, where patient alleles are absent in the donor, drive GvHD risk through donor T-cell recognition of recipient antigens [11]. Conversely, HvG mismatches, where donor alleles are absent in the patient, influence graft rejection risk through recipient immune responses against donor cells. Our analysis reveals that these directional mismatches frequently differ, with approximately 30% of mismatched patient-donor pairs having a difference between HvG and GvH.

Additionally, traditional matching algorithms compute mismatches per locus and then sum across loci, potentially overestimating total incompatibility. For instance, a donor-patient pair with one GvH mismatch at HLA-A and one HvG mismatch at HLA-B would be classified as having two mismatches by traditional algorithms, despite never having more than one mismatch in either direction across the entire genotype. GRIMM-II’s approach of computing the maximum of total GvH and total HvG mismatches across all loci more accurately represents the biological reality and a large number of potentially suitable donors in the two- and three-mismatch categories. This expanded donor pool is particularly valuable for patients from underrepresented ethnic minorities who face substantially lower probabilities of finding perfectly matched donors [14, 8]. The graph-theoretic framework underlying GRIMM-II provides several advantages beyond computational efficiency [23]. The modular architecture naturally accommodates the dynamic nature of donor registries, enabling incremental updates as new donors are added without requiring complete recomputation. The separation of genotypes into class I and class II components, with systematic enumeration of subclass vertices representing genotypes missing one allele, creates a searchable structure that dramatically reduces the candidate space while guaranteeing that no valid matches are missed.

For imputation, ML-GRIM’s two-stage approach addresses the combinatorial explosion that occurs with nine-locus typing. By performing classical GRIM imputation on a reduced set of three loci followed by consistency checking against remaining typed loci, the algorithm maintains accuracy while avoiding the prohibitive memory requirements of simultaneously tracking all possible phasing combinations across nine loci [17, 16]. This approach proves particularly effective given the linkage disequilibrium structure of HLA, where information from highly polymorphic loci like A, B, and DRB1 substantially constrains possibilities at other loci. Our analysis of mutual information between loci highlights that selection of informative loci for typing significantly impacts imputation accuracy, and suggests optimal loci to type when not all loci can be typed.

Several limitations of the current implementation warrant consideration. First, while the algorithm efficiently handles up to three mismatches, computational performance degrades when extended to larger mismatch thresholds such as those encountered in haploidentical transplantation [12]. Alternative algorithmic approaches may be required for scenarios involving 4-5 mismatches, though the clinical utility of automated matching in such heavily mismatched scenarios remains uncertain since in practice, there are practically no patients who do not have a large number of 3-mismatched or less donor.

Second, the current implementation treats all mismatches equivalently, not accounting for emerging evidence regarding permissive versus nonpermissive mismatches based on structural epitope differences [7]. Integration of HLA epitope matching algorithms, particularly for DPB1 T-cell epitope groups, represents a logical extension that would enhance clinical applicability and will be implemented next. Similarly, incorporation of population-specific mismatch tolerance data, if such information becomes available from outcomes research, could enable more nuanced donor ranking. Importantly, the ML-GRMA framework can be easily adapted to include differences in mismatches.

## Acknowledgments

LG was funded by NIAID R01AI173095 and NIDDK R01DK139240

## 7 Appendix

### 7.1 Donor candidates claims

***Claim:*** Two classes that have one mismatch between them share 1 subclass. ***Proof:*** The subclass where the unmatched allele was removed is the same for both classes.

To prove the next claim, we will use the claim above.

***Claim:*** Every two genotypes that share 7 or more alleles must also share at least one subclass.

***Proof:*** If the genotypes share 10 alleles, all their subclasses match. If the genotypes share 9 alleles, there is one class that matches, and also all of its subclasses match, and another class with one mismatch, which is one matched subclass. If they share 8 alleles, if both alleles are in the same class, there is a match in the other class, therefore a match in all its subclasses. Otherwise, there is one mismatch in each class, which is one matched subclass for each class. If they share 7 alleles, if all 3 alleles are in the same class, all the subclasses of the other class match. Otherwise, there is one class with one mismatch, and another with two mismatches. The class with the one mismatch has one shared subclass. ***Definition:* Candidates** are genotypes from the Donor’s Graph that match a genotype from the Patient’s Graph with more than 7 matches.

### 7.2 Python vs. Cython

To efficiently handle large datasets and computationally intensive tasks related to genotype analysis, we implemented several key functions using Cython, a superset of Python designed to give C-like performance with code written mostly in Python. These optimizations include functions for filtering and comparing genetic data. The code includes filters out genotypes with less than 7 matches by comparing patient genotypes against multiple donor genotypes to count allele matches. The Hash table provides hashing genotypes and transforms them into compact, consistent values that enable fast lookups and comparisons in large databases. This allows for quick filtering and retrieval of relevant genotypes and efficient similarity counting without repeatedly processing entire sequences, ensuring scalability and speed in largescale data processing tasks.

### 7.3 Validation datasets and methods

**Table 1:**
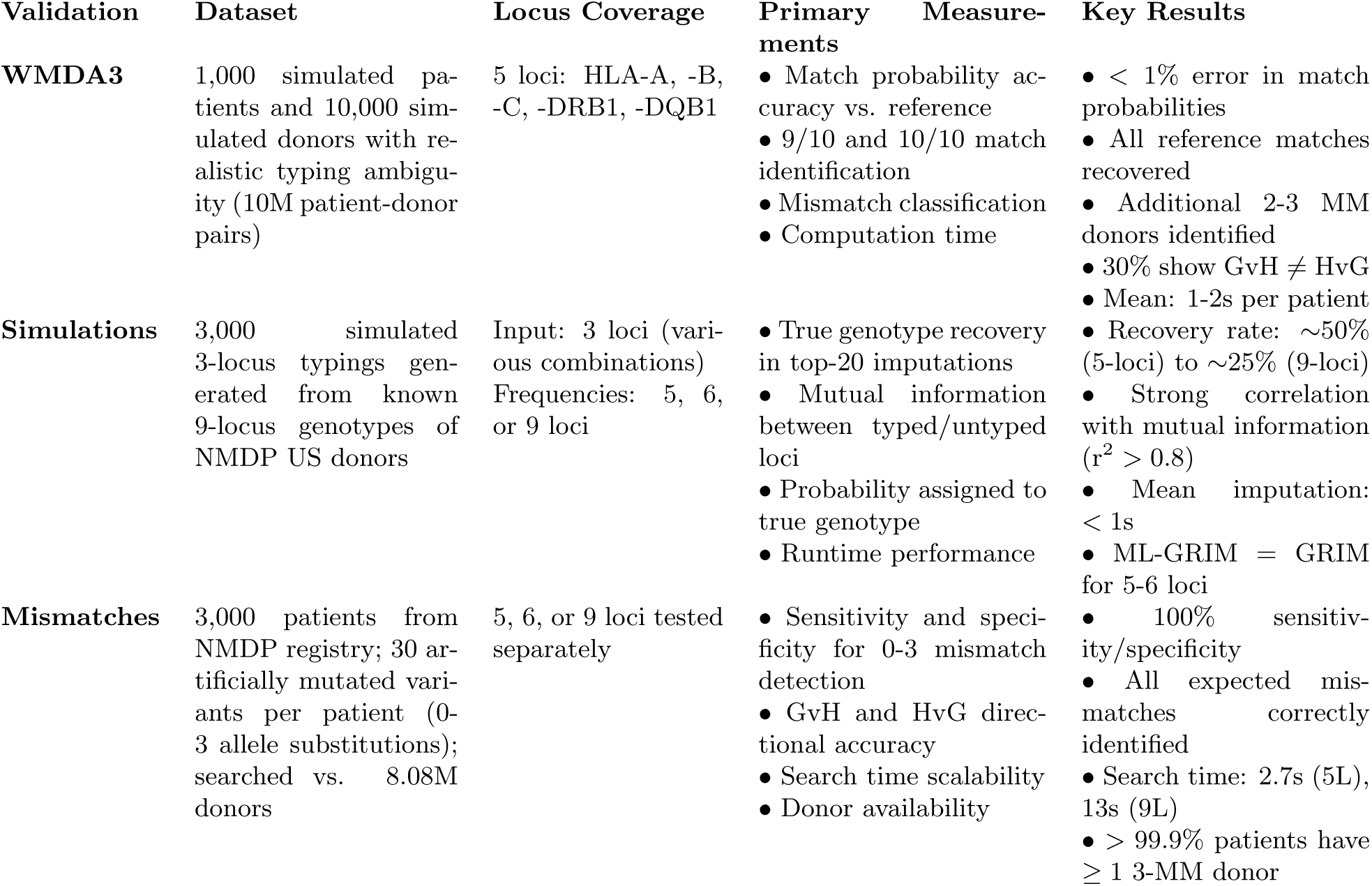
Summary of validation approaches for ML-GRIM and ML-GRMA algorithms. Three complementary validation strategies were employed to assess imputation accuracy, matching sensitivity/specificity, and probabilistic match score computation. MM = mismatch; GvH = graft-versus-host; HvG = host-versus-graft; L = loci; s = seconds, Simulations = Simulted typings with different combinations of 3 loci, Mismatches = Simulated mismatches

